# Deciphering the role of mucosal immune responses and cervicovaginal microbiome in resistance to HIV infection in HIV-exposed seronegative (HESN) women

**DOI:** 10.1101/2021.05.07.443078

**Authors:** Sivasankaran Munusamy Ponnan, Kannan Thiruvengadam, Chaitanya Tellapragada, Anoop T Ambikan, Aswathy Narayanan, Sujitha Kathirvel, Janani Shankar, Akshaya Rajaraman, Mehar Afsan Amanulla, Thongadi Ramesh Dinesha, Selvamuthu Poongulali, Shanmugam Saravanan, Kailapuri Gangatharan Murugavel, Soumya Swaminathan, Vijayakumar Velu, Barbara Shacklett, Ujjwal Neogi, Luke Elizabeth Hanna

## Abstract

The female genital tract (FGT) is an essential site of HIV infection. Discerning the nature of HIV-specific local immune responses is crucial for identifying correlates of protection in HIV-exposed seronegative (HESN) individuals. The present study involved a comprehensive analysis of soluble immune mediators, secretory immunoglobulins (sIg) and levels of natural killer (NK) cells, CXCR5^+^ CD8^+^T cells, T follicular helper cells (Tfh) and T regulatory cells (T regs) in the vaginal mucosa, as well as the nature and composition of the cervicovaginal microbiome in HESN women. We found significantly elevated antiviral cytokines, soluble immunoglobulins, and increased frequencies of activated NK cells, CXCR5^+^ CD8^+^ T cells and Tfh cells in HESN females as compared to HIV unexposed healthy (UH) women. Analysis of the genital microbiome of HESN women revealed a greater bacterial diversity and increased abundance of *Gardnerella spp* in the mucosa of HESN women. The findings suggest the female genital tract of HESN females represents a microenvironment equipped with innate immune factors, antiviral mediators and critical T cells subsets that protect against HIV infection.

## Introduction

Mucosal surfaces are primarily protected by innate immune mechanisms, but adaptive immune mechanisms also operate (1). The natural barriers present in the female genital tract (FGT) are insufficient to protect against all infections, though the FGT is a site of immunological balance (2). Innate immune cells play a vital role in the non-specific destruction of foreign organisms. Natural Killer (NK) cells recognize infected host cells via surface receptors and employ various mechanisms to kill them (3). Increased NK cell activity is reflected by the higher cytotoxic capacity of NK cells and increased production of NK cell-specific cytokines and chemokines associated with control of infection (4-6). Perturbations in the NK cell receptor repertoire, including cells expressing natural cytotoxicity receptors (NCR) and killer inhibitory receptors (KIR), have been reported in HIV and other viral infections (7). Elevated NK cell activity has been correlated with protection against infection in several high-risk cohorts of HIV-exposed seronegative (HESN) subjects, including intravenous drug users, HIV-1 discordant couples and perinatally exposed infants (8). Studying the role of KIR and NCR expressing NK cell subsets may help us understand NK cell-mediated control of early HIV infection.

Neutralization has long been viewed as the primary effector function of humoral immunity and is considered to be the primary correlate of antibody-mediated protection in HIV infection (9). However, the role of mucosal antibodies in HIV infection continues to be controversial (10). Soderlund et al. (2007) detected HIV-1-specific neutralizing IgA antibodies in some infected persons, but these antibodies were unable to block the transfer of the virus from dendritic cells (DC) to susceptible target cells (11). On the other hand, Nag et al. found that HIV-1 gp120-specific IgG in the cervical fluid could mediate ADCC activity and that their levels correlated inversely with genital viral load (12).

Antibody responses are mainly dependent on CD4^+^ T cell help, and the significant subset that provides this help is the T follicular helper (Tfh) cell. These cells help in B cell affinity maturation, isotype switching, and the development of long-lived plasma cells (13). A high frequency of peripheral blood Tfh cells has been reported to correlate with high titers of broadly neutralizing antibodies and reduced viral load in HIV-infected individuals (14). Similarly, infiltration of CD8^+^ T cells into the B cell follicle area has been reported in HIV-infected individuals. More recently, Shen et al. demonstrated that CXCR5^+^ CD8^+^ T cells residing in the germinal center follicular area play an essential role in neutralizing virus-infected target cells (15). However, the role of CXCR5^+^ CD8^+^ T cells or Tfh cells in disease control in highly exposed uninfected individuals, and the correlation of these cell types with HIV-specific antibody titers is yet to be assessed.

The cervicovaginal microbiome is a complex ecosystem with a preponderance of *Lactobacillus* spp. Vaginal *lactobacilli* prevent the colonization of the female genital tract by pathogenic microbes by maintaining an acidic pH. In addition, the cervicovaginal mucus functions as a physical barrier for the ascent of exogenous pathogens in women with a healthy vaginal microbiome (16). Depletion of vaginal *lactobacilli* and cervicovaginal mucus is commonly observed among women with conditions such as bacterial vaginosis, aerobic vaginitis, genital chlamydiosis, and Human Papilloma Virus infection (17). Many studies have reported cervicovaginal dysbiosis as an essential risk factor for the acquisition of HIV-1 infection (16, 18, 19). Cervicovaginal colonization by bacterial species such as *Prevotellabivia, Sneathia* spp. and *Mycoplasma hominis* have been reported to be associated with increased concentrations of pro-inflammatory cytokines and chemokines in the FGT and enhanced susceptibility to HIV-1 acquisition (20-22).

The present study attempted to characterize in detail the role of mucosal NK cells, CXCR5^+^ CD8^+^ T cells, Tfh, soluble immune mediators, antiviral cytokines, and chemokines, as well as the composition of the cervicovaginal microbiome in the early control of infection in HIV-exposed seronegative women.

## Results

### Sociodemographic features of the study population

There was no significant difference between the two groups for age, practice of vaginal douching, presence of bacterial vaginitis, *Chlamydia trachomatis* (CT), or *Neisseria gonorrhoeae* (NG) infection. All study participants were screened for cervical cancer by testing for Human papillomavirus (HPV) infection using the Pap smear test. The HESN group reported sex on an average of 2 times a month with their infected partner, with or without condom use.The mucosal specimens collected for this study were obtained at least a week after the last sexual act and precisely two weeks after the start of the last menstrual period in the volunteers (Table-1).

**Table1:** Demographic and clinical characteristics of HESN and HIV Unexposed Healthy women.

| Parameters |  | HESN women<br>(N = 37) | Unexposed<br>women (N = 35) |
| --- | --- | --- | --- |
| Age | Median | 36 | 33 |
|  | Range | 27-42 | 22-42 |
| Vaginal douching |  | 37 (100%) | 35 (100%) |
| Cervical Cancer (HPV) Positive |  | 0 | 0 |
| STD (BV, CT, NG) Positive |  | 0 | 0 |
| Average number of sexual contacts per month | With condom (median) | 2 | 2 |
|  | Without condom (median) | 1 | 1 |
|  | Unknown (median) | 1 | 0 |
| HIV-infected partner on ART | Yes | 37 (100%) | 0 |
|  | No | 0 | 35 (100%) |
| HESN-HIV-1 exposed seronegative; HPV-Human Papilloma Virus; BV-Bacterial vaginitis; CT-*Chlamydia trachomatis*; NG- *Neisseria gonorrhoeae*. |  |  |  |

### Presence of increased frequency of mucosal CD8^+^ T cells expressing follicular homing receptor in HESN individuals

Studies have shown that the frequency of CD8^+^ T cells expressing the follicular homing receptor, CXCR5, correlates inversely with viral load in both human and animal models (15, 23). A recent study reported that elite controllers had increased numbers of CXCR5^+^ CD8^+^ T cells exhibiting increased effector function, and suggested that these cells have a potential role in the natural control of HIV infection (24). Similarly, it is well known that follicular T helper cells play an important role in class switching and antibody production. However, the distribution of Tfh in the mucosal microenvironment has not been fully elucidated. Hence, we analyzed the frequencies of CXCR5^+^CD8^+^ T cells, Tfh, and Tregs in cytobrush specimens obtained from HESN women and compared these values with those detected in unexposed healthy (UH) women.

To examine thefrequency and distribution of follicular homing/CXCR5+ CD8+ T cells, Tfh cells and Tregs, we stained the cervical cells obtained by cytobrush sampling of the cervix of HESN and UH women with fluorochrome tagged monoclonal antibodies and performed multicolor flow cytometry (Table S1). As previously described, CD4^+^ Tfh cells were defined as CD4+ CD45RO+ CCR7+ PD1+ CXCR5+ CXCR3-cells (14) and follicular homing CD8^+^ T cells were defined as CD8+ CD45RO+ CCR7+ PD1+ CXCR5+ CXCR3-cells (25). Treg cells were defined as CD3+ CD4+ CD127-CD25+ cells (Fig. 1a).

**Figure 1:**
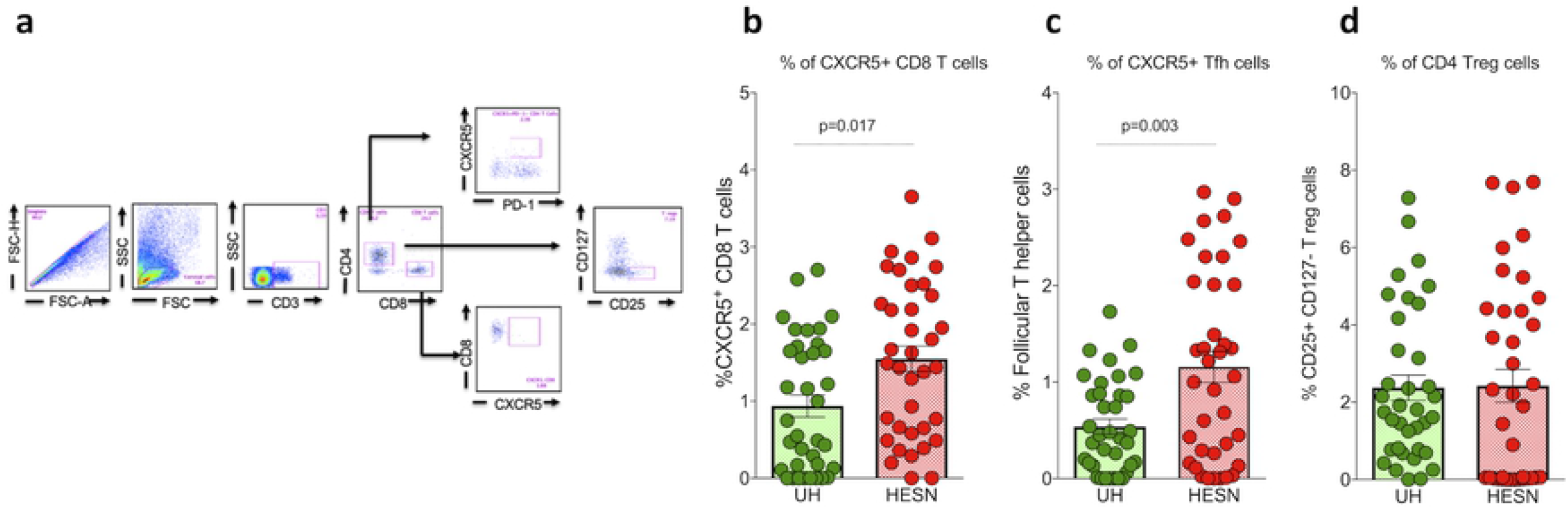
Mucosal CXCR5^+^CD8+ T cells, T follicular helper cells and Tregs: **(a)** Representative flow plots showing the % frequency of CXCR5^+^ T cells and Treg cells in the study groups (HESN -N=37, and HU -N=35). **(b)** Figure showing the cumulative frequency of CXCR5^+^ CD8+ T cells in the HESN and UH groups. **(c)** Figure showing the cumulative frequency of T follicular helper cells in the HESN and UH groups. **(d)** Figure showing the cumulative frequency of CD127-CD25+ T reg cells in the HESN and UH groups. The scatter dot plots summarize the % frequency of total CD8^+^CXCR5^+^ T cells, Tfh and Treg cells (median, 1^st^, and 3^rd^ quartiles). P values were determined using the Mann-Whitney test.

We found that the HESN group had significantly increased numbers of mucosal follicular homing CXCR5^+^ CD8^+^ T cells (p=0.017) (Fig.1b) as compared to the UH group. We further analyzed the expression of CXCR5 on CD4^+^ Tfh cells and found that the frequency of CXCR5 expressing CD4^+^ Tfh cells was significantly higher in the HESN group as compared to the UH group (p=0.003)(Fig. 1c). The median frequency of mucosal CXCR5^+^ CD8^+^ T cells was 1.49% (range: 0.65 -2.37%), and the median frequency of mucosal Tfh cells among the CD4^+^ T cell subset was 1.06% (range: 0.26 -2.01%) in the HESN group as compared to 0.44% (range: 0.14 -0.87%) in the UH group. On the other hand, there was no difference in the distribution of mucosal T regulatory cells between the HESN and UH groups (Fig. 1d).

### Abundance of natural cytotoxicity receptor (NCR) expressing NK cells in the cervical mucosa of HESN individuals

NK cell-mediated antiviral effector function and ADCC mediated by NK cells and antiviral antibodies have been implicated in the control of HIV infection (26, 27). This evidence suggests a role for NK cells in the control of HIV and opens up a new avenue for research in the development of HIV vaccines (28). However, NK cell function and distribution in the mucosal tissue are not fully understood. We analyzed the distribution of NK cells expressing NCR and KIR in the genital mucosa of HESN women and UH women using multicolour flow cytometry on cells present in the cervical cytobrush specimens. NK cells expressing activating natural cytotoxicity receptors (Table S2) were defined as CD3-CD16+ CD56+ CD27+ NKG2D+ Nkp30+/Nkp44+/Nkp46+ cells, and NK cells expressing inhibitory killer cell immunoglobulin-like receptors were defined as CD3-CD16+ CD56+ CD27+ KLRG1+ CD158a+/CD158b+/CD158e1+ cells as described by Kulkarni et al. (5) (Figure S1& S2).

We found significantly higher frequencies of NK cells expressing NCRs such as Nkp30, Nkp44 and Nkp46 in the genital mucosa of HESN women as compared to UH women. We performed Boolean analysis of NCR-expressing cells to identify multiple receptor expressing cells and found that dual NCR-expressing NK cells (Nkp30+Nkp44+, Nkp44+Nkp46+ and Nkp30+Nkp46+) were also significantly elevated in the HESN group (Fig 2a). In contrast, the frequency of NK cells expressing inhibitory receptors were similar in the HESN and UH groups. However, CD158b+ and CD158a+CD158e1+ dual receptor-expressing NK cells alone were found to be significantly higher in the genital mucosa of HESN women as compared to UH women (P=0.018 and p=0.031, respectively) (Fig 2b).

**Figure 2:**
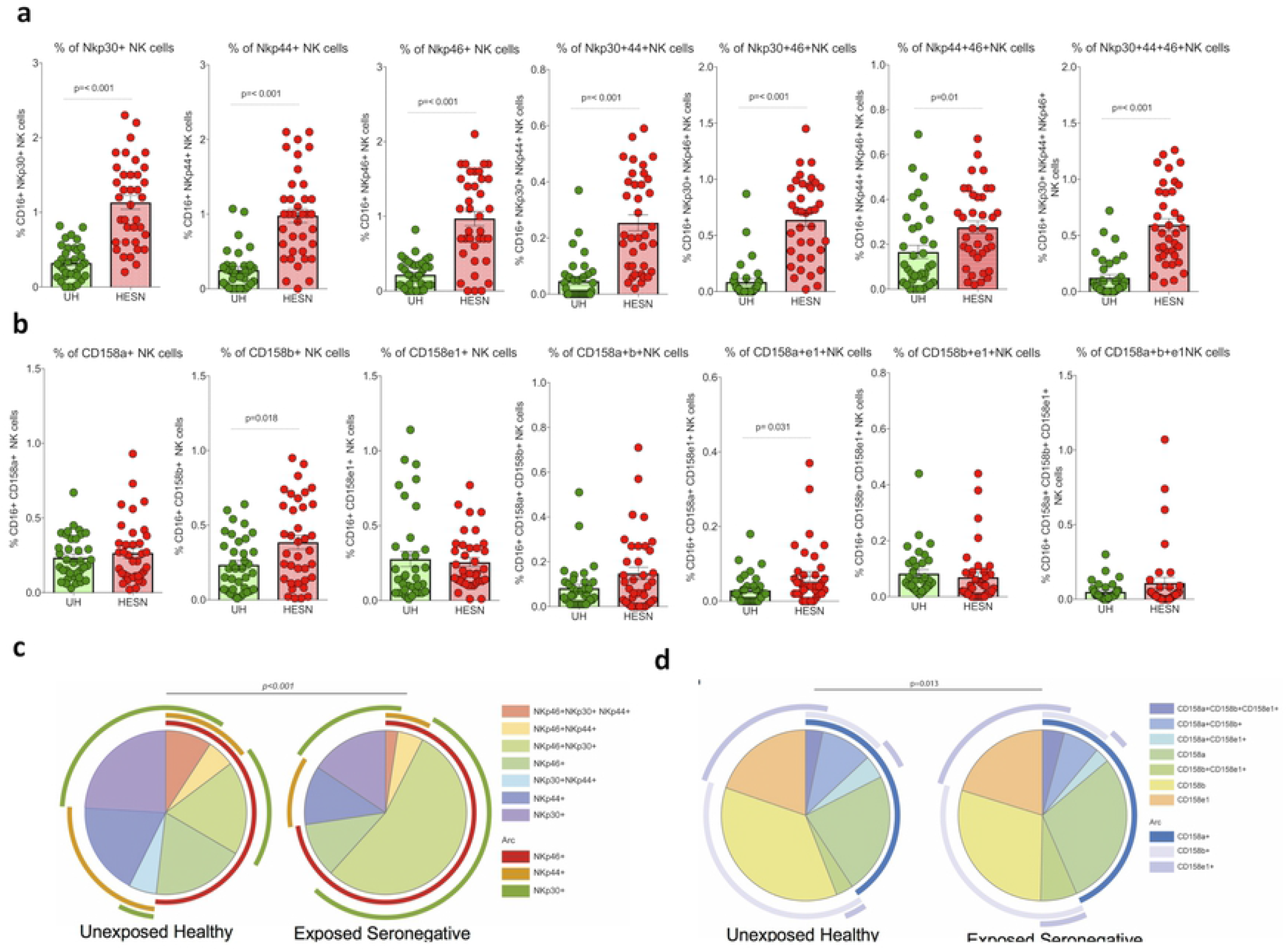
Distribution of natural cytotoxicity receptor (NCR) and Killer cell immunoglobulin-like inhibitory receptor (KIR) co-expressing mucosal NK cells: **(a)** Proportion of NK cell expressing each activating receptor (Nkp30, Nkp44, Nkp46) **(b)** Proportion of NK cell expressing each inhibitory receptor (CD158a, CD158b, CD158e1). The scatter dot plots summarize the % frequency of total NCR and KIR expressing NK cells from cytobrush specimens (HESN -N=37, and UH -N=35). The graphical plots show the median, 25^th^, and 75^th^ percentiles and range (IQR). P values were determined using the Mann-Whitney test. **(d)**SPICE analysis of different combinations of natural cytotoxicity receptor (NCR) expressing NK cells indicated statistically significant differences in the permutation of different combinations. **(e)** SPICE analysis of the different combination of Killer cell immunoglobulin-like inhibitory receptor (KIR) expressing NK cells indicated statistically significant differences in the permutation of different combinations.

We also found that NK cells expressing all 3 NCRs (Nkp30+Nkp44+Nkp46+) were significantly more abundant in the HESN group. However, there was no difference in the frequency of NK cells expressing multiple KIR (CD158a+CD158b+CD158e1+ cells) between the HESN and UH groups (p>0.950). These data taken together, demonstrate that NK cells expressing natural cytotoxicity receptors (NCR) are more abundant in the mucosa of HESN women as compared to UH women (**Fig. 2c, d**).

### Increased levels of total secretory IgG and IgA in the mucosal microenvironment of HESN individuals

Mucosal immunoglobulin (IgG, IgA and IgM) levels were determined in the study participants’ cervicovaginal lavage using a BD cytometric bead array. We found increased levels of total IgG and IgA in HESN women as compared to UH women. The median level of mucosal IgG was 10,000 ng/ml (2874-55454 ng/ml) in the HESN group and 454 ng/ml (101 -3516 ng/ml) in the UH group. The median level of mucosal IgA was 6447 ng/ml (1984 - 11442 ng/ml) in the HESN group and 1955 ng/ml (487 - 7351 ng/ml) in the UH group. Thus, it is evident that the HESN participants had significantly higher levels of total IgG and IgA in their cervicovaginal secretions than UH women (IgGp<0.001 and IgAp=0.024, respectively); however, there was no significant difference in IgM levels between the two groups (Fig. 3).

**Figure 3:**
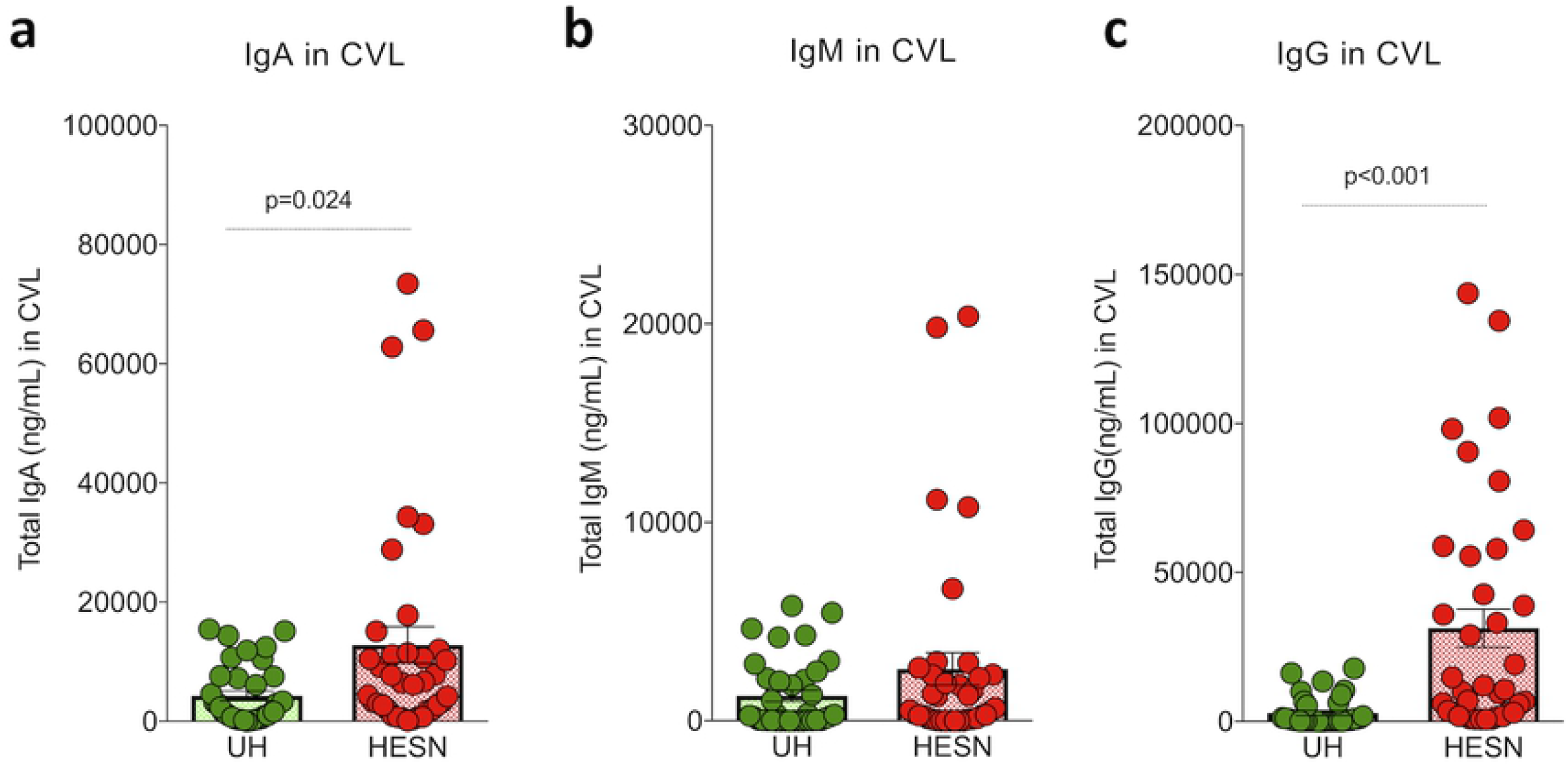
Levels of mucosal immunoglobulin (IgG, IgA, and IgM) levels in cervical vaginal lavage: Mucosal immunoglobulin (IgG, IgA, and IgM) levels in CVL specimens measured by cytometric bead array and reported in ng/ml. **(a)** Total IgA level in cervicovaginal lavage **(b)** Total IgM level in cervicovaginal lavage **(c)** Total IgG level in cervicovaginal lavage. The box-and-whisker plots represent median, 25^th^, and 75^th^ percentiles and range (IQR). P values were determined using the Mann-Whitney test. (Note: HESN - HIV exposed seronegative, (N=37): UH - HIV unexposed seronegative (N=35)).

### Differences in dynamics of pro-inflammatory molecules, antiviral cytokines and chemokines in the cervicovaginal secretion of HESN individuals

Earlier studies that reported elevated antiviral responses against HIV in HIV controllers, slow progressors and HIV exposed seronegative individuals (28, 29), have demonstrated a significant role for cytokines and chemokines in HIV disease progression (30). As part of our analysis, we measured antiviral cytokines and chemokines in the cervicovaginal lavage of HESN and UH individuals using the Cytometric Bead Assay (LEGENDplex™).

Significantly elevated levels of several T cell/NK cell cytokines and soluble factors like IL-2, IFN-γ, TNF-α, IL-17α, IL-17f, IL-21, IL-4, IL-5, IL-9, IL-10, IL-13, and Granzyme B were detected in the HESN group (Fig. 4b & 4c). The HESN cohort also had significantly higher levels of antiviral cytokines like IFN-α2, IFN-β, IFN-λ2/3, IL-12p70 and GM-CSF (Fig. 4a). Increased levels of few more cytokines were found in the HESN cohort, but the increase was not significant (Table S4). In contrast, levels of all the pro-inflammatory chemokines measured were found to significantly lower in the HESN group. These included IL-8, IP-10, Eotaxin, TARC, MCP-1, RANTES, MIP-1α, MIP-1β, MIG, ENA-78, MIP-3α, GROa and I-TAC (Fig. 4d; Table S5). These findings suggest that the mucosal microenvironment in HESN individuals is more quiescent and has a good supply of antiviral effector cytokines and chemokines which makes it unsuitable for establishment of HIV infection.

**Figure 4.**
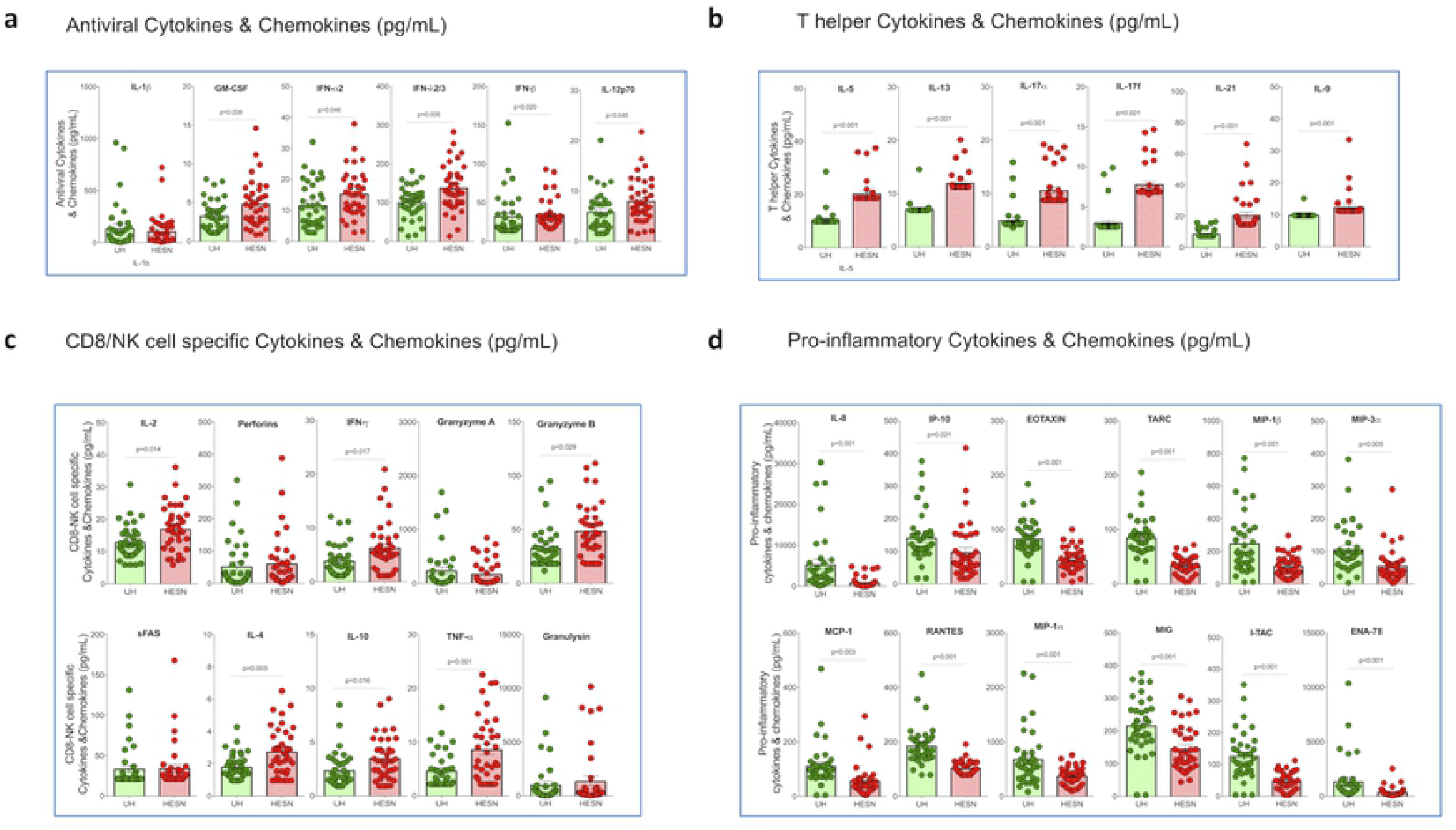
Levels of anti-viral mediators, T helper cell-specific and CD8^+^ T cell/NK cell-specific effector molecules, cytokines and chemokines in the cervicovaginal lavage: Elevated levels of **(a)** antiviral molecules **(b)** T helper cell cytokines (IL-1β, IP-10, IFN-λ1, IL-12p70, IFN-α2, IFN-λ2/3, GM-CSF, IFN-β, IL-10), **(c)** CD8/NK cell specific cytokines (IL-2, IL-4, IL-10, IL-6, IL-17, TNF-α, sFAS, sFASL, IFN-γ, granzyme A, granzyme B, perforin, granulysin) cytokines and chemokines and **(d)** Pro-inflammatory chemokines (IL-8, IP-10, EOTAXIN, TARC, MCP-1, RANTES, MIP-1α, MIG, ENA-78, MIP-3α, GROa, I-TAC, MIP-1β) in the CVL of HESN (N=37) and UH (N=35) groups. The data shown is the median level of cytokines and chemokines (pg/mL). The scatter dot plots show the median, 25^th^, and 75^th^ percentiles and range (IQR). P values were determined using the Mann-Whitney test. (Note: HESN -HIV exposed seronegative; UH -HIV unexposed seronegative controls).

### Composition of the cervicovaginal microbiome differs between the HESN and UH groups

Microbiome analysis revealed that bacteria belonging to the phyla Proteobacteria, Epsilonbacteria, Acidobacteria, and Chloroflexi were relatively more abundant (p <0.001) in the HESN group as compared to the UH group (Fig. 5a). The family level relative abundance of the microflora in the two groups is presented in Fig.5b. Group-wise comparison of the microbiome composition at the genus level was performed using PERMANOVA, which indicated that *Gardnerella* had the highest PERMANOVA coefficient (Fig.5c). Our findings are in agreement with previous observations of higher abundance of *Gardnerella* and Intermediate abundance of *Lactobacillus spp* in highly exposed commercial sex workers from the Nairobi cohort (31). Microbiome distribution at the genus level is shown in Fig. 5d & 5e. At the species level, we found that majority (51%) of the UH women had a preponderance of *L. iners* (cervicotype-2 flora), followed by *Gardnerella vaginalis* (CT-3 flora) in 29% and mixed anaerobic bacteria (CT-4 flora) in 11% (Fig. 6a). Among the HESN group, CT2, CT3 and CT4 flora were observed in 32, 47 and 11% of the women respectively. The comprehensive cervicotype classification system was adapted from Anahtar et al., 2015 (21). Interestingly, our study subjects did not include women with the low-risk CT1 cervicotype. Taken together, our findings reveal significantly higher microbial diversity (abnormal flora belonging to CT-3 and CT-4) in the HESN cohort (58%) as compared to the control group (40%) (Fig.6a). Statistically significant differences were detected between the study groups in the relative abundance of *Veillonella montpellierensis, L. gasseri, L. crispatus, Prevotellaamnii, Sneathiaamnii* and *Streptococcus agalactiae* (Fig 6b). While *Veillonella montpellierensis, L. gasseri and L. crispatus* were more abundant in the HESN group, *Prevotellaamnii, Sneathiaamnii* and *Streptococcus agalactiae* were more common in the UH group (Fig 6b). Higher abundance of ungrouped bacteria was also observed in the HESN group as compared to the UH group in our cohort.

**Figure 5:**
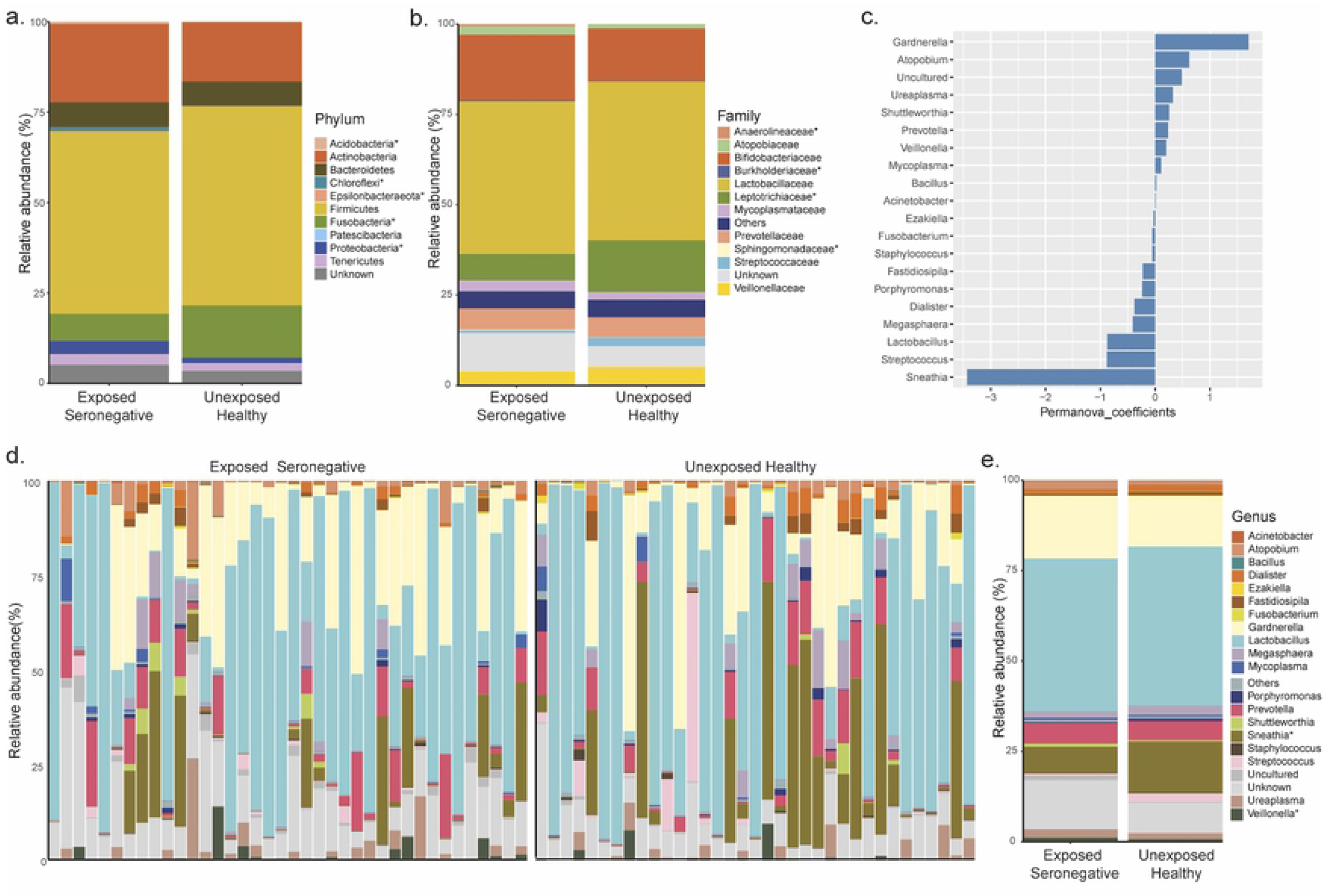
Composition of the cervicovaginal microbiome in the study population. **(a)** Mean relative abundance in the phylum **(b)** Mean relative abundance in Family. **(c)** Top 20 taxa at genus level which have a significant effect on separating the groups as per PERMANOVA analysis **(d)** Visualization of sample-wise relative abundance (%) in the two experimental groups **(e)** Mean relative abundance in Genus.* denotes significance in the Mann-Whitney test.

**Figure 6:**
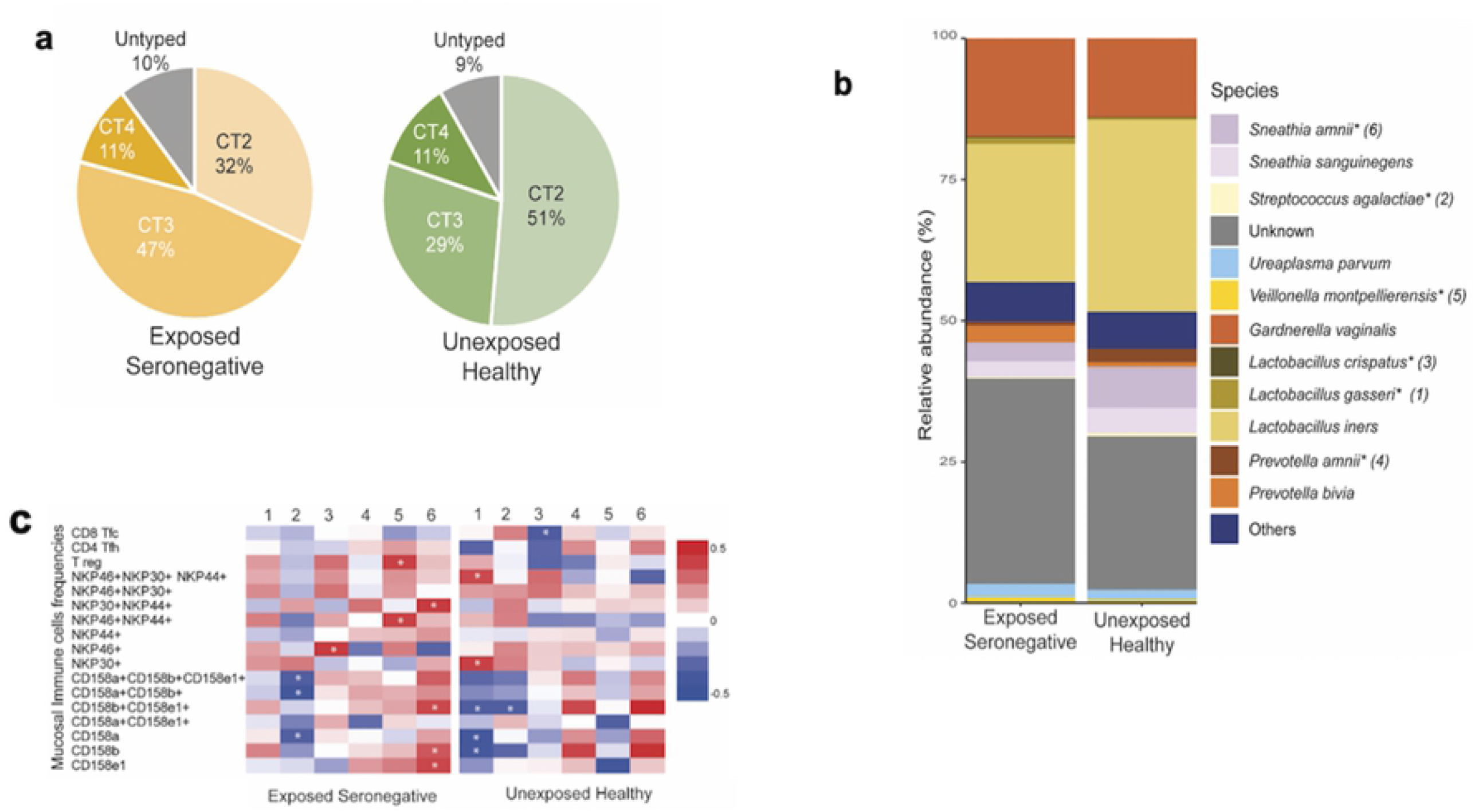
Distribution of bacterial species and their association with inflammation. **(a)** Distribution of CT2, CT3 and CT4 bacteria in HESN and UH women. **(b)** Bar graph showing average relative abundance (%) of various species. * denotes significance in the Mann-Whitney test. **(c)** Heat map showing Spearman’s correlation between significant species (from Mann-Whitney test) and mucosal immune cell frequencies. * denotes significant correlation.

Interestingly, we observed a positive correlation between the frequency of mucosal Tregs and NK cells and higher microbial diversity in the HESN group, suggesting that increased colonization of the genital mucosa with diverse microbial species might lead to increased recruitment of critical components of the innate immune system that coordinate the early control of HIV infection. The correlation observed between bacterial species and immunological markers in the study groups is shown in Fig. 6c. The intricate role of mucosal immune cell activation in determining the pattern and diversity of the local microbiome remains incompletely understood. Understanding the interplay between the microbiome and the innate and adaptive immune system may shed light on some of the critical events that orchestrate protection against HIV infection.

## Discussion

Mucosal immune responses present at the surfaces that serve as points of contact with the pathogen are believed to play a critical role in controlling/preventing HIV infection (32). Innate immune responses are the first line of defense against microbes (33). Earlier studies have documented that host factors present in the FGT, including the natural microbiota, play a role in both protection and susceptibility to infection (1, 34). Protective mucosal responses against HIV are best characterized in individuals who remain relatively resistant to HIV infection despite repeated exposure (35). This phenomenon has been documented in many groups, including commercial sex workers (36), serodiscordant couples (37), infants born to HIV-infected mothers, and others. Long-term non-progressors (LTNPs) constitute another group of individuals who are HIV infected but relatively resistant to disease progression. These individuals sometimes share common immunological features with HIV controllers and have been used to understand the correlates of resistance/susceptibility to HIV infection and disease progression (1, 37, 38).

Several correlates of protection against HIV infection have been previously described in both systemic and genital compartments. HIV-specific CD8^+^ T cell responses (39) and the presence of anti-HIV IgA (40) have been implicated as markers of protection or indicators of exposure (41). Increased levels of the molecules RANTES, serpin B, alpha-2 macroglobulin-like molecule and cystatin have been correlated with resistance to infection (42). Although multiple correlates of resistance to infection and/disease progression have been described, the mechanisms through which the different factors operate have not been deciphered. Many future investigations are required to understand the mechanisms involved in the protection against HIV-1 infection.

Characterizing the subset of innate immune cells like the NK cells present in FGT may help us to understand the nature of the innate immune mechanisms operating in the mucosal microenvironment and provide a clue to the development of an adaptive immune process. An earlier study reported elevated levels of NK cell activity and higher secretion of NK cell-specific cytokines in highly exposed uninfected intravenous drug users compared to controls (43). In addition to the evidence supporting the role of NK cells and elevated levels of IFN-γ in mediating control of HIV infection, changes in expression of different cell surface receptors on NK cells have also been implicated in HIV infection and control (44).

Stimulation of NK cell cytotoxicity involves interaction between target cell antigens and stimulatory surface receptors on NK cells (5, 45-47). Wang et al. reported that elevated expression of NCR correlated with low viremia and that NKp30 plays a significant role in NK cell activation, degranulation and cytotoxicity (4). Another study reported that increased expression of activating receptors and potency of cytotoxic NK cells correlated with reduced viral load in recently infected individuals and long term non-progressors (5). Our study also found significantly higher levels of NK cells expressing NKp46, NKp44 and NKp30 in the HESN cohort as compared to the UH population. We further analyzed frequencies of dual receptor and triple receptor-expressing NK cells using Boolean gating and found that the proportion of activated NK cells expressing multiple NCR was also significantly higher in the HESN cohort. This observation supports the hypothesis that the activating receptor-expressing NK cell subset might have a role to play in the early control of HIV infection. We also examined the expression of inhibitory receptors (CD158a, CD158b, CD158e1 and NKG2A) on NK cells and found that the proportion of cells expressing these receptors was correspondingly lower in the HESN group as compared to the UH group. Alter et al (2009) also suggested that activating KIR, or weakly inhibitory KIR, may have a significant role to play in viral infections (7). Zwolinska et al. found that the frequency of NK cells expressing KIR receptors was elevated during HIV disease progression, and suggested a role for these cells in regulating the exaggerated innate and adaptive antiviral response (48). Studies done in African female sex workers (FSW) have shown decreased genetic pairing of inhibitory KIR with its HLA ligand, leading to a decrease in inhibitory signalling. These observations support a role for NK cells in protecting against virus transmission and demonstrate that improved NK activation or reduced NK inhibition is linked with resistance to HIV infection (49, 50).

High levels of inflammatory cytokines and tissue damage in the FGT have been suggested to increase susceptibility to HIV infection (51). Earlier studies have documented increased inflammation in the FGT of healthy donors as compared to the periphery. These observations suggest that decreased levels of pro-inflammatory mediators in the FGT would lower inflammation of the mucosa and reduce the risk of HIV acquisition. The findings of our study provide evidence for decreased levels of pro-inflammatory mediators in the mucosa of HESN women as compared to UH women, as well as higher levels of antiviral molecules such as Type I and III interferons and cytotoxic effector molecules, suggesting that mechanisms are in place to control inflammation of the FGT in HESN women, while responding to low-level viral replication. Lajoie et al. showed that the mucosal immune system of FSWs is less activated as compared to that of low-risk HIV-negative women; they also found that this group of individuals had much lower mucosal levels of pro-inflammatory cytokines/chemokines compared to sexually active non-FSWs (52). These findings indicate that low levels of mucosal immune activation triggers the local innate response without increasing the number of potential target cells (52). Our study also found lower levels of pro-inflammatory cytokines in the mucosal compartment of HESN women as compared to UH individuals. Lower levels of pro-inflammatory molecules in the mucosa possibly contributes to the integrity of the genital mucosa by controlling inflammation and maintaining tight junctions, thus favoring the control of HIV infection. It is, therefore, crucial to consider the genital tract as a separate entity when studying HIV-1 transmission and evaluating HIV resistance.

Few studies in HESN individuals have shown that these individuals have neutralizing IgA antibodies in the genital mucosa, which could protect against infection (53). Some reports have suggested that the presence of anti-HIV IgA antibodies in genital secretions correlates more strongly with exposure than resistance to HIV infection (54). In the present study, we measured total IgG, IgA and IgM in the CVL specimen of HESN and UH cohorts and observed elevated levels of IgA and IgG in the HESN group as compared to the UH group. However, there was no difference in the level of IgM between the two groups. Though the study did not measure HIV-specific antibodies, we hypothesize that the abundance of antibodies at the mucosal surface of HESN women may contribute to protection against HIV infection, perhaps by the indirect killing of HIV-infected cells through antibody-dependent cell-mediated cytotoxicity coordinated by NK cells (55).

Antibody responses typically require CD4^+^ T cell help, and the major helper subset that provides this help is the T follicular helper (Tfh) cell (56) that primes the B cells to produce antibodies. In HIV-infected individuals, a high frequency of circulating Tfh cells has been found to correlate with high titers of broadly neutralizing antibodies (bNAb) and reduced viral load (14, 57). Sivasankaran et al. reported that a subtype C specific HIV vaccine induced circulating memory-like Tfh cells that correlated positively with variable region specific antibody production (58). More recent studies have shown that CXCR5^+^ CD8^+^ T cells traffic into the B cell follicle area of SIV-infected rhesus macaques and suppress residual viral replication (59). In the present study, we evaluated the frequencies of mucosal CD4^+^ Tfh and CXCR5^+^ CD8^+^ T cells in cytobrush specimens and found elevated levels of these T cell subsets in HESN women as compared to UH women. To our knowledge, this is the first study reporting higher frequencies of Tfh, CXCR5^+^ CD8^+^ T cells in HESN cohorts from India. These results suggest that the induction of mucosal CD4^+^ Tfh cells and CXCR5^+^ CD8^+^ T cells might contribute to the early control of infection, possibly through induction of a significant antibody response or some other unknown mechanism to eradicate infected cells at the mucosal level.

HIV infection results in the production of pro-inflammatory cytokines, and their levels increase with the progression of HIV disease (60). Many soluble mediators can suppress HIV replication *in vitro* (61). The number of cytokines produced and cytokine receptor density on the host cell surface are believed to be key determinants for the control of HIV infection (60). Many innate soluble markers possess an inhibitory effect on HIV, including α, β, θ defensins, IL-6, IL-10, IL-16, TNF-α, interleukin 1β, and type I interferons α and β (62). In the present study, we analyzed several antiviral cytokines and chemokines in the CVL of HESN and UH women using cytometric bead array and observed significantly elevated levels of IL-12p70, IFN-α2, IFN-λ2/3, GM-CSF, IFN-β and IL-10 in the HESN group. We also analyzed cytokines produced by CD8 and NK cells among others and found that IL-2, IL-4, IL-10, TNF-α, IFN-γ and granzyme B were present at significantly increased levels in HESN women. Besides, sFAS, sFASL and perforin levels were also higher in the mucosal compartment of HESN women, though the difference between groups was not significant. Furthermore, we analyzed the levels of Th cytokines and found that IL-5, IL-13, IL-9, IL-17α, IL-17f, IL-21 and IL-22 were very high in HESN women. The presence of high levels of immune cell-specific cytokines and antiviral cytokines and chemokines at the mucosal surface is thought to be instrumental in instructing the development of different types of immune mechanisms that contribute to the early control of HIV infection.

The cervicovaginal microbiome is a dynamic ecosystem, and the diversity of its composition can vary due to multiple factors, including ethnicity, socio-economic status, level of education, sexual behavior, and lifestyle factors (18). An interesting finding from the present study concerning the cervicovaginal microbiome was a higher relative abundance of *L. iners* in both groups as compared to *L. crispatus*, unlike in other studies (21, 22, 63). Reasons for the low abundance of *L. crispatus*in our cohort could not be assessed due to the lack of epidemiological data (socio-economic status, level of education and lifestyle factors) regarding the vaginal health status (by Gram-stained smear examination) of the study participants. Low et al. reported that intravaginal practices, as well as the presence of BV, can increase the risk of HIV acquisition. However, intravaginal cleaning with soap, disruption of vaginal flora, and usage of vaginal microbicides can reduce the likelihood of HIV acquisition (19). In a recent study on South-African women, it was reported that women with a predominance of *L. crispatus* were more resistant to the acquisition of HIV as compared to those with a predominance of *L. iners* and other anaerobic bacterial flora. The investigators also reported an increase in pro-inflammatory cytokines among women colonized with bacteria such as *Vellionella montpellierensis, Prevotellabivia* and *Sneathia sanguinegens* (64). However, in the present study, we observed a predominance of *L. Iners* and high level of diversity in the cervicovaginal microbiome of the HESN individuals. Studies report that a lactobacillus-dominant vaginal microbiota generally reflects vaginal health and is associated with a decreased risk of HIV acquisition (16, 64). We observed higher abundance of *L. Crispatus* in the HESN group and a low level of pro-inflammatory mediators that may help maintain the mucosal surface’s integrity and reduce the risk of HIV acquisition in these individuals. Further, we observed a correlation between mucosal NCR-expressing, activated NK cells and mucosal microbiota in HESN women, suggesting that immune cell trafficking and activation of immune cells in the cervicovaginal surfaces may reduce the risk of HIV infection in association with vaginal dysbiosis.

To summarize, the majority of HIV infections occur via the vaginal or rectal route, and therefore, many researchers believe that a strong pre-existing immune response in the mucosa-associated lymphoid tissue (MALT) may be able to protect against HIV infection. In addition to antigen-specific adaptive immune responses which were not evaluated in this study, there is mounting evidence that factors associated with inflammation, or alternatively, immune quiescence, may contribute to HIV susceptibility at the mucosal surface (29, 39). The present study explored some of the immune mechanisms operating in the mucosal microenvironment of HESN women and identified mucosal natural cytotoxicity receptor-expressing NK cells, CXCR5^+^ CD8^+^ T cells, follicular T helper cells, and soluble markers, as well as a highly diverse cervicovaginal microbiome, to be potential correlates of protection against HIV-1 infection. However, further investigation is necessary to determine the functional capabilities of these factors in abrogating HIV infection. The multifactorial nature of HIV resistance emphasizes the importance of the interplay between immune cells and the composition of the vaginal microbiome at the mucosal level. Further understanding in this line would help us to devise strategies to enhance useful innate and adaptive immune responses to prevent HIV infection. This knowledge would also prove to be useful for the identification of novel candidate vaccines.

## Materials and Methods

### Ethics statement

The study protocol was approved by the Scientific Advisory Committee of the ICMR-National Institute for Research in Tuberculosis (NIRT), Chennai, India. The study was conducted under Good Clinical Laboratory Practice (GCLP) guidelines. The study protocol was reviewed and approved by the Institutional Ethics Committee of ICMR-NIRT (IEC ID-2015015) and the Institutional Review Board of Y. R. Gaitonde Centre for AIDS Research and Education (YRG CARE; YRG-302), Chennai, India.

### Study participants

The study participants consisted of two groups of individuals: seronegative female spouses of HIV-1 seropositive men (HIV discordant couples) (HESN, n=37) and HIV-unexposed seronegative women (UH, n=35). Enrolment in the study required willingness to provide written informed consent for specimen collection and storage. HIV-exposed seronegative women aged between 20-35 years with a history of multiple unprotected sex events with an HIV infected partner during the past one year were recruited into the HESN group, and HIV negative healthy women aged between 20-35 years with no history of HIV exposure were recruited into the control group. Women outside the reproductive age group, women on menstruation, those with STIs or other major illnesses, moribund individuals as well as pregnant and lactating women were excluded from the study.

### Sample collection

#### Collection and processing of cervical cytobrush specimens

Mucosal specimens were collected by a trained physician. At least two cytobrush specimens were obtained from each study participant. The cytobrush was inserted just within the cervical os and rotated to one 360° turn. Samples contaminated visibly with blood were discarded. Immediately after sampling, the cytobrush was placed in a 15 mL tube containing 3 mL of RPMI 1640 medium containing 100 U/mL penicillin, 100 µg/mL streptomycin, and 2.5 µg/mL amphotericin B, and placed on ice. The specimens were transported on ice to the laboratory and processed within 3 hours of collection. The cytobrush was gently rotated several times in the transport medium to release all the cells into the medium and then discarded. The cell suspension was centrifuged at 330g for 10 minutes, and the pellet was resuspended in complete RPMI medium containing 100 U/mL penicillin and 100 µg/mL streptomycin and used for flow cytometric analyses.

#### Collection and processing of cervicovaginal lavage (CVL)

CVL was collected by gently washing the cervicovaginal area with 10 mL of sterile normal saline (pH, 7.2) and withdrawing the fluid using a 5 mL syringe. Following CVL collection, samples were frozen as quickly as possible at -80°C. At the time of analysis, samples were thawed at room temperature, centrifuged at 10,000×g for 5 minutes, and supernatants were analyzed for soluble antiviral and innate immune factors.

### Quantification of total mucosal IgG, IgA and IgM antibodies

Total IgG, IgA and IgM antibodies were quantified in the CVL specimens using the human immunoglobulin cytokine bead array (CBA) flex set assay (BD Biosciences, San Jose, CA, USA). The Ig CBA assay was performed as per the manufacturer’s instructions. Briefly, 50 µl of sample, 1:2 diluted standards (1:2, 1:4, 1:8, 1:16, 1:32, 1:64, 1:128, 1:256) and negative control (plain media) were added to the respective tubes. About 50 µl of capture bead was added to each tube and incubated for 1 hour at room temperature. The tubes were washed with 1 mL of wash buffer at 200g for 5 minutes, and the supernatant was removed. 50 µl of PE detection reagent was added to the pelleted beads and incubated for 2 hours at room temperature. The washing procedure was repeated, and the pellet was resuspended in 300 µl of wash buffer and mixed well. Samples were acquired on a FACS ARIA III SORP flow cytometer (BD Biosciences), and the FCS files were analyzed using the FCAP array v3 software (BD Biosciences).

### Measurement of soluble markers in CVL specimens using cytometric bead array

Pro-inflammatory chemokines including MCP-1 (CCL2), RANTES (CCL5), IP-10 (CXCL10), Eotaxin (CCL11), TARC (CCL17), MIP-1α (CCL3), MIP-1β (CCL4), MIG (CXCL9), MIP-3α (CCL20), ENA-78 (CXCL5), GROα (CXCL1), I-TAC (CXCL11) and IL-8 (CXCL8), CD8^+^ T cell/NK cell cytokines and cytolytic molecules including IL-2, IL-4, IL-10, IL-6, IL-17, TNF-α, sFAS, sFASL, IFN-γ, granzyme A, granzyme B, perforin andgranulysin, antiviral cytokines including IL-1β, IL-6, IL-8, IL-10, IL-12p70, IFN-α, IFN-β, IFN-λ1, IL-29, IFN-λ2/3, IL-28, IFN-γ, TNF-α, IP-10 and GM-CSF, and Th cell cytokines including IL-5, IL-13, IL-2, IL-6, IL-9, IL-10, IFN-γ, TNF-α, IL-17A, IL-17F, IL-4, IL-21 and IL-22 were measured in CVL using the Bio legend LEGENDplex human multianalyte kit based cytometric bead array (CBA) (Bio legend, CA, USA) following manufacturer’s recommendations. Briefly, beads coated with 13 specific capture antibodies were mixed. Subsequently, 50 μL of the mixed capture beads, 50μL of CVL diluted 1:2 or 1:20, and 50 μL of detector antibody was added and incubated for 2 hours on a plate shaker (250 rpm) at room temperature in the dark. 50 μl of streptavidin-phycoerythrin (SA-PE) detection reagent was then added to each assay tube and incubated for 30 minutes on a plate shaker (approximately 250 rpm) at room temperature in the dark. The samples were washed with 1 mL of wash buffer (at 200 g) for 5 minutes. The bead pellet was resuspended in 100 μL of wash buffer after discarding the supernatant. Samples were analyzed on a BD FACS ARIA III SORP flow cytometer and analyzed using the LEGENDplex data analysis software, v8.0 (Biolegend, CA, USA). Individual cytokine concentrations were indicated by their fluorescence intensities. Cytokine standards were serially diluted to facilitate the construction of calibration curves, which were necessary for determining the protein concentration of the test samples (65). The lower limit of detection of each molecule is provided in Supplementary Table S3.

### Characterization of immune cell types in the cervical cytobrush specimen

Mucosal cytobrush-derived cells were washed twice, and viability was evaluated by using the trypan blue dye exclusion method. The average percentage of viability was in the range of 80%. The cells were stained with the following cocktail of monoclonal antibodies for enumeration of the different immune cell types (Table S1 & S2).

#### T follicular helper and Treg panel

CD3-APC H7 (SK7-BD Bioscience), CD4-PERCP Cy5.5 (RPA-T4-BD Bioscience), CD8-BUV 737 (RPA-T8-BD Bioscience), CD45RO BUV395 (UCHL-1-BD Bioscience), CCR7-PEcy7 (G043H7-BD Bioscience), CXCR3-APC R 700 (IC6-BD Bioscience), CXCR5-BB515 (RF8B2-BD Bioscience), PD-1-PE (EH12.1-BD Bioscience), CD25-APC (M-A251-BD Bioscience) and CD127-PECF594 (HIL.7R.M21-BD Bioscience).

#### NCR expressing NK cells

CD3-APC H7 (SK7-BD Bioscience), CD16-BUV737 (3G8-BD Bioscience), CD56-APC Alexa 700 (NCAM-BioLegend), CD27-BB515 (MT271-BD Bioscience), NKG2D-PECF594 (ID11-BD Bioscience), NKP44-PE (p44-8-BD Bioscience), NKP46-PECY7 (9E2-BD Bioscience), NKP30-APC (p30-15-BD Bioscience).

#### KIR expressing NK cells

CD3-APC H7 (SK7-BD Bioscience), CD16-BUV737 (3G8-BD Bioscience), CD56-BUV396 (NCAM 16.2-BD Bioscience), CD27-PECY7 (MT271-Bio Legend), KLRG1-PECF94 (231A2-BioLegend), CD158a-PE (CH-L-BD Bioscience), CD158b-BB515 (CH-L-BD Bioscience), CD158e1-APC (DX9-BD Bioscience).

The cells were stained for 20 minutes at 4°C. About 0.5 million cells were stained for each panel. After staining, the cells were washed, fixed with BD Cytofix (2% paraformaldehyde), and analyzed on a FACS ARIA III SORP flow cytometer (Becton Dickinson) (66). A minimum of 300,000 total events was acquired for each panel, and data were analyzed using FlowJo software, version 10.5.4 (Tree Star Inc., Ashland, Oregon, USA).

### Analysis of the vaginal microbiome by next-generation sequencing (NGS)

The V3-V4 region of 16s rRNA was sequenced using Illumina HiSeq2500 with 2×250 cycles chemistry. The raw reads were checked for base quality, base composition, and GC content using the FastQC tool (version 0.11.8). After the quality check, pre-processing steps were employed on the data. Firstly, forward V3 and reverse V4 primer sequences were trimmed using an in-house PERL script. Properly paired paired-end reads with Phred quality score >20 were considered for V3-V4 consensus generation. The pre-processed reads were then analyzed using QIIME (67) software (version 1.9.1) to pick up OTUs with 97% similarity cut-off and taxonomy classification with the SILVA database as reference (68). Various R packages were used for all downstream statistical analyses and figure generation. Mann-Whitney test was performed at phylum, family, genus and species level between the two groups using R package stats since the data followed a non-normal distribution. PERMANOVA significance test was performed at the genus level using R package vegan (69). Correlation analysis was performed between relative abundance at the species level, immune cell frequencies, and soluble markers by Spearman correlation using R package psych (70).

### Statistical analysis and graphical representation of the soluble factors and immunological markers

The statistical analysis and graphical representation of immunological markers were carried out using the data analysis program SPICE (version 6.0) (71) and ggplot2 ver 3.2.0 (https://ggplot2.tidyverse.org/). Statistical analyses of mucosal immune cells and soluble markers were performed using GraphPad Prism, version 7.05 (GraphPad Software, Inc., CA). Values are presented as median, interquartile range, and percentage. Mann-Whitney test was used to examine the difference in frequency (%) of different immune cell subsets and levels of soluble markers between the two groups. Correlation analysis was performed to determine the relationship between the frequency of different immune cell types and levels of mucosal soluble markers. For all analyses, differences were considered significant if the p-value was <0.05.

## Author Contributions

S.M.P., L.E.H. and S.S. designed the conceptual framework of the study, designed experiments, and wrote the paper. S.M.P performed the experiments and analyzed the data. K.T. contributed to statistical analyses. T.R.D., S.S., S.P. and K.G.M contributed to specimen collection. C.T., A.T.A., A.N. and U.N contributed to the vaginal microbiome analysis by next-generation sequencing. S.P.T., S.S., A.R., J.S., S.K., V.V. and B.S contributed to the review and editing of the manuscript. All authors provided valuable input throughout the study.

## Supplementary Information

**Figure S1 -Representative pseudocolor FACS plot of activating natural cytotoxicity receptor (NCR) expressing NK cells**: Mucosal NCR expressing NK cells were enumerated in cytobrush specimens of HESN and UH participants. (a) CD3-ve lymphocytes were first gated, then CD16+ and CD56+ NK cells and CD27+ NKG2D+ cells were gated, followed by gating on NKP30+, NKP44+, NKP46+ cells using Boolean gate to identify multiple NCR receptor-expressing NK cells. (b) Representative ancestry and back gating strategy used for identification of NKP 44+ activated NK cells.

**Figure S2 -Representative pseudocolor FACS plot of killer cell immunoglobulin-like inhibitory receptor (KIR) expressing NK cells**: Mucosal KIR expressing NK cells were enumerated in cytobrush specimens of HESN and UH participants. (a) CD3-ve lymphocytes were gated, then CD16+ and CD56+ NK cells and CD27+ KLRG-1+ cells were gated, followed by gating on CD158a+, CD158b+, CD158e1+ cells using Boolean gate to identify multiple KIR expressing NK cells. (b) Representative ancestry and back gating strategies used for the identification of CD158e1+ activated NK cells.

**Supp. Table 1:** Commercial reagents used for Tfh cell and Treg immune-phenotyping by multicolor flow cytometry.

**Supp. Table 2:** Commercial reagents used for NK cell immune-phenotyping by multicolor flow cytometry.

**Supp. Table 3:** The theoretical limits of detection of cytokines and chemokines by cytometric bead array.

**Supp. Table 4:** Antiviral cytokines, chemokines and T helper cells cytokines in cervicovaginal lavage.

**Supp. Table 5:** CD8+ T cell/NK cell cytokines and cytolytic molecules in cervicovaginal lavage.

